# A high throughput multi-locus insecticide resistance marker panel for tracking resistance emergence and spread in *Anopheles gambiae*

**DOI:** 10.1101/592279

**Authors:** Eric R. Lucas, Kirk A. Rockett, Amy Lynd, John Essandoh, Nelson Grisales, Brigid Kemei, Harun Njoroge, Christina Hubbart, Emily J. Rippon, John Morgan, Arjen Van’t Hof, Eric O. Ochomo, Dominic P. Kwiatkowski, David Weetman, Martin J. Donnelly

## Abstract

The spread of resistance to insecticides in the mosquito vectors of diseases such as malaria and dengue poses a threat to the effectiveness of control programmes, which rely largely on insecticide-based interventions. Monitoring the resistance status of mosquito populations is therefore essential, but obtaining direct phenotypic measurements of resistance is laborious and error-prone. In contrast, high-throughput genotyping offers the prospect of quick and repeatable estimates of resistance, while also allowing the genotypic markers of resistance to be tracked and studied. We developed a panel of 28 known or putative markers of resistance in the major malaria vector *Anopheles gambiae*, which we use to test the association of these markers with resistance and to study their geographic distribution. We screened resistance-phenotyped *An*. *gambiae* from populations from a wide swathe of Sub-Saharan Africa (Burkina Faso, Ghana, Democratic Republic of Congo (DRC) and Kenya), and found evidence of resistance association for four mutations, including a novel mutation in the detoxification gene *Gste2* (*Gste2*-119V). We also identified a gene duplication in *Gste2* which combines a resistance-associated mutant form of the gene with its wild-type counterpart, potentially alleviating the costs of resistance. Finally, we describe the distribution of the multiple evolutionary origins of *kdr* resistance, finding unprecedented levels of diversity in the DRC. This panel represents the first step towards developing a quantitative predictive genotypic model of insecticide resistance that can be used to screen *An*. *gambiae* populations and predict resistance status.

## 1 Introduction

It is estimated that the use of insecticide-treated nets and indoor residual spraying of insecticide has been responsible for 78% of 663 million cases of malaria averted in the period 2000-2015 (Bhatt et al. 2015). The documented rise and spread of resistance to currently-used insecticides in the major malaria mosquito vectors therefore presents a worrying trend that may negatively impact malaria control programmes (Implications of Insecticide Resistance Consortium 2018, Kafy et al. 2017, Protopopoff et al. 2018). In particular, the major malaria vector in Sub-Saharan Africa (SSA), *Anopheles gambiae*, now has widespread resistance to pyrethroids, and is showing pockets of high resistance to carbamates and organophosphates (Ranson & Lissenden 2016). To maintain effective malaria control, accurate assessment of the insecticide resistance profile of mosquito populations is essential. Furthermore, as new insecticides are brought to market, it will be crucial to identify and track resistance to these new products in order to react appropriately before the failure of malaria control.

Phenotypic assessment of insecticide resistance requires live mosquitoes of fixed age to be assayed in carefully controlled conditions. The effort and care required to perform these assays make them difficult to perform on a large scale and prone to inconsistency if conditions are not rigorously controlled (Kleinschmidt et al. 2018). Moreover, phenotyping approaches are only sensitive when resistance has reached appreciable frequency in the population. For these reasons, there is great interest in the prospect of screening mosquitoes for the genetic signatures of insecticide resistance, which would provide a convenient and simple way of assessing resistance from any type of mosquito collection method, including dead mosquitoes routinely collected by monitoring and evaluation programmes. Genetic screening carries the additional benefit of improving our understanding of the origins of resistance mutations, which can inform resistance management policies. For example, by understanding where resistance originates, research can begin to elucidate the environmental and population-level contexts which favour its rise and spread. This has been seen in the evolution of drug resistance in the malaria parasite, which repeatedly appears and spreads from an area in Cambodia (Miotto et al. 2013), driving research into understanding the causes of this pattern.

There are two main challenges involved in the development of genetic screening assays for insecticide resistance. First, genetic markers of resistance need to be identified and their effect size estimated. Several single nucleotide polymorphism (SNP) mutations associated with insecticide resistance have already been discovered in *Anopheles* (Du et al. 2005, Jones et al. 2012, Martinez-Torres et al. 1998, Mitchell et al. 2014, Ranson et al. 2000, Riveron et al. 2014, Weetman et al. 2018, Weill et al. 2003), providing the first opportunities for genetic screening. However, a substantial proportion of the variance in resistance remains unexplained, making it crucial to identify new markers and to quantify their impact.

Second, new approaches are required for large-scale, high-throughput screening of large marker panels, to reduce the costs and time required to screen mosquito populations. The Agena Biosciences iPLEX MassARRAY technology is a platform capable of accurate genotyping combined with high level multiplexing capability. MassARRAY panels have already been developed for a range of infectious disease questions, such as studying resistance to malaria in humans (Malaria Genomic Epidemiology Network et al. 2014), mapping genetic diversity in parasites (Volkman et al. 2007) and detecting ancestry and inter-specific hybridisation in *Anopheles* (Norris et al. 2015)

Here, we present the development and application of an iPLEX MassARRAY panel of 28 genotypic markers for screening *Anopheles gambiae* populations to quantify the effects of known and putative insecticide resistance markers, and to track the origins of spread of knockdown resistance variants (*kdr*) in the Voltage-gates sodium channel (*Vgsc*) gene. We applied our panel to samples from four different countries from West (Burkina Faso and Ghana), Central (Democratic Republic of Congo - DRC) and East Africa (Kenya). These samples were assayed for susceptibility to permethrin (all four countries), deltamethrin (Ghana, DRC and Kenya) and DDT (Ghana). This genetic screen:

1. revealed a heterologous duplication in *Gste2* that combines resistance-associated and wild-type alleles,
2. identified an unprecedented diversity of *kdr* genetic backgrounds in the DRC,
3. reported the first evidence of a role for the *Gste2*-119V mutation in insecticide resistance in *An*. *gambiae*.
4. found evidence that a mutation found alongside *kdr* reduces insecticide resistance, possibly indicating that this mutation serves to compensate for physiological costs of *kdr*.

## 2 Methods

### 2.1 Sample collection

#### 2.1.1 Samples from Burkina Faso

*Anopheles* larvae were collected using hand dippers from semi-permanent and temporary water bodies between June and October 2014 at four sites (Bakaridjan: 10.407N, −4.562W; Bounouba: 10.357N, −4.439W; Naniagara: 10.536N, −4.669W and Tiefora: 10.632N, −4.556W). Samples were transported to the insectaries in Banfora where they were maintained at a temperature of 27°C (±2°C) and a relative humidity of 80% (±10%), and fed with TetraMin Baby®. Adult mosquitoes were tested for insecticide resistance using the CDC bottle bioassay (Brogdon & McAllister 1998) with 20ppm permethrin. All mosquitoes were stored individually in perforated PCR tubes and placed into sealed bags with silica gel to avoid the decomposition and ensure DNA preservation. DNA was extracted using the LIVAK method (Livak 1984) and samples were identified to species using SINE PCR (Santolamazza et al. 2008).

#### 2.1.2 Samples from DRC

Samples from the DRC were the same as those reported by Lynd et al. (2018). *An*. *gambiae* mosquitoes were collected in March and April 2016 from three rural collection sites (Pambwa, 3.937 N, 20.772 E; Fiwa, 4.318 N, 20.778 E; Bassa, 4.267 N, 21.283 E) in the area surrounding the major town of Gbadolite, near the border with the Central African Republic. Adult mosquito collections were carried out in all three villages using both manual and mechanical (‘Prokopack’, Vazquez-Prokopec et al. 2009) aspirators and were maintained in a field insectary until egg-laying. Larvae were collected from breeding sites in Fiwa and Pambwa. Larvae collected directly, and those raised from eggs, were reared until the adult stage. All mosquitoes were identified to species group using phenotypic keys (Gillies & Coetzee 1987) and insecticide resistance testing was carried out on 3-5 day-old adult *An*. *gambiae s*.*l*. to assess resistance to permethrin (0.75%) and deltamethrin (0.05%) using standard World Health Organization (WHO) protocols, and Abbott’s correction (World Health Organization 2016). All mosquitoes were stored on silica gel in 0.2ml tubes for later DNA analyses. DNA was extracted from individual mosquitoes using Nexttec (Nexttec, Biotechnologie GmbH) extraction plates according to manufacturer’s instructions, and mosquitoes were identified to species using SINE PCR (Santolamazza et al. 2008).

#### 2.1.3 Samples from Ghana

Ghanaian samples were collected from Keta (5.917 N, 0.991 E), an urban community in the Volta Region of Ghana. Keta is located within the coastal savannah agroclimatic zone, with vegetation consisting of shrubs, grasses and a few scattered trees.

*An*. *gambiae* s.l. larvae were collected from September to November, 2016, using hand-held ladles. Collections were made from a variety of habitats such as pools, puddles, drainage channels, irrigations and vegetable fields, and transported to the laboratory in partly-filled labelled plastic containers. Larvae were reared in the insectaries at the Animal Science department of the Biotechnology and Nuclear Agricultural Research Institute (BNARI), Accra. Insecticide resistance assays were performed on 3-5 day-old females following standard WHO tube assay protocols.

DNA was extracted from single mosquitoes using the Nexttec 96-well plate DNA Isolation kit. Species identification was performed on the whole-genome-amplified DNA (Section 2.2) using standard species identification PCR (Scott et al. 1993) followed by SINE PCR to further distinguish *An*. *gambiae* from *An*. *coluzzii* (Santolamazza et al. 2008).

#### 2.1.4 Samples from Kenya

Mosquitoes were sampled from four malaria-endemic regions in Western Kenya (Teso: 0.656 N, 34.353 E, Bondo: −0.098 S, 34.273 E, Rachuonyo: −0.353 S, 34.656 E and Nyando: −0.162 S, 34.921 E). These regions were under distinct vector control interventions, with Teso and Bondo having insecticide treated nets and Nyando and Rachuonyo having both insecticide treated nets and indoor residual spraying. *Anopheles* larvae were collected from aquatic habitats using the standard dipping method and transferred to plastic tins using wide-mouth pipette for transportation to KEMRI, Kisumu laboratories for rearing. Larvae were reared on a mixture of fish food and brewer’s yeast provided daily. Upon pupation, individuals were transferred to cages to emerge as adults and provided with 10% sucrose in cotton pledgets.

Three-day-old adult females were exposed to either permethrin (0.75%) or deltamethrin (0.05%) using WHO impregnated papers for 1 hour following WHO guidelines (World Health Organization 2013). Mortality was recorded and all mosquitoes were placed in individual tubes and frozen at −20°C for molecular analysis. DNA was extracted from whole sample using ethanol precipitation (Collins et al. 1987).

A newly-developed melt-curve based species identification assay (Chabi et al. 2018) was used because existing techniques are prone to interpretation errors as a result of similar sized PCR bands for *An*. *arabiensis* vs *An*. *gambiae* on agarose gels. The assay uses SYBR-green with a universal forward primer (5’-ATTGCTACCACCAAAATACATGAAA-3), a reverse primer matching both *An*. *arabiensis* and *An*. *gambiae* with G-8 extension (5’-GGGGGGGGGAATAATAAGGAACTGCATTTAAT-3’) to slightly increase the amplicon melting temperature, and an *An*. *arabiensis* specific reverse primer (5’-GGATGTCTAATAGTCTCAATAGATG-3’).

### 2.2 SNP genotyping

We developed a multiplex panel of 28 SNP markers (Supplementary Data S1) based on the AgamP3 reference genome (Sharakhova et al. 2007) and information on variable sites identified by the *An*. *gambiae* 1000 Genomes (Ag1000G) project (*Anopheles gambiae* 1000 Genomes Consortium 2017). Primers for these SNPs were developed using the MassARRAY Assay Design software (version 4.0.0.2; Agena Biosciences, Hambrug, Germany).

The SNPs that we chose fell into three categories. First, we included eight SNPs that have previously been associated with insecticide resistance. These include the two *kdr* mutations (*Vgsc*-995F and *Vgsc*-995S), as well as a further *Vgsc* mutation (*Vgsc*-1570Y) known to exist on a *Vgsc*-995F background and conferring increased resistance to pyrethroids (Jones et al. 2012), two mutations associated with resistance to dieldrin (*Rdl*-296G and *Rdl*-296S, Du et al. 2005), a mutation in the gene *Gste2* associated with metabolic resistance to DDT (*Gste2*- 114T, Mitchell et al. 2014) and a mutation associated with resistance to carbamates and organophosphates (*Ace1*-280S, previously referred to as 119S, Weill et al. 2003). The eighth SNP in this set was a mutation in codon 119 of *Gste2* (*Gste2*-119V), which was included because a mutation in the same codon has been shown to strongly increase resistance to DDT in *An*. *funestus* (Riveron et al. 2014), raising the prospect that the mutation in *An*. *gambiae* may have a similar function.

Second, we included eight non-synonymous SNPs (*Vgsc*-1868T, *Vgsc*-1874S, *Vgsc*-1874L, *Vgsc*-1853I, *Vgsc*-1934V, *Vgsc*-1746S, *Vgsc*-791M and *Vgsc*-1597G) that have been found to be strongly associated with the *Vgsc*-995F mutation (*Anopheles gambiae* 1000 Genomes Consortium 2017) and are thus regarded as potential candidates influencing resistance either by enhancing the impact of *Vgsc*-995F or compensating for suspected fitness effects.

Third, we included twelve SNPs that can differentiate the five haplotype backgrounds of the *Vgsc*-995F mutation (haplotype backgrounds F1-F5) and the five haplotype backgrounds of the *Vgsc*-995S mutation (haplotype backgrounds S1-S5) (Anopheles gambiae 1000 Genomes Consortium 2017). These SNPs were identified using the Ag1000G data by searching for SNPs that differentiated each haplotype background from the other haplotypes with the same *kdr* mutation. Within each of the two *kdr* mutations (*Vgsc*-995F and *Vgsc*-995S), the SNPs always provided perfect separation between the five possible haplotype backgrounds, although a few haplotypes that were not assigned any background in the Ag1000G data could be incorrectly assigned a background based on these SNPs (Supplementary Data S2).

DNA extractions were whole-genome amplified and then genotyped with the Agena Biosciences iPLEX platform as described (Fabrigar et al. 2016), using our panel of 28 SNPs. Genotype calls were obtained from the raw data using the TYPER software (version 4, Agena Biosciences) and raw data as cluster plots were inspected and manually curated to remove ambiguous calls. To assess the quality of the assays and the genotype calling process, we also used our panel to genotype 287 samples previously used in phase 2 of Ag1000G (https://www.malariagen.net/data/ag1000g-phase-2-ar1) and compared the two sets of genotype calls. Raw data and genotype calls are provided in Supplementary Data S3, and genotype calls are also provided along with sample information in Supplementary Data S4. Samples that failed genotyping at more than 50% of loci were excluded from the analysis (this filter removed 39 samples from Ghana and two samples from Kenya).

### 2.3 Deviations from Hardy-Weinberg equilibrium

Tests for deviations from Hardy-Weinberg equilibrium were performed using the *HWExact* function from the R package *HardyWeinberg*. These tests were performed for every SNP x location combination in which the SNP was segregating (i.e.: where both alleles were present in the sampling location). This resulted in 72 separate tests, which were adjusted for multiple testing using a Bonferroni correction, with a resulting threshold α value of 0.0007.

### 2.4 Identification of kdr haplotype backgrounds

The process of identifying the *kdr* haplotype background of a sample based on its genotype was implemented using custom R functions that has been made available as supplementary materials and which is described in Supplementary Methods S1.

### 2.5 Statistical analysis of resistance

The two SNPs in *kdr* (*Vgsc*-995F and *Vgsc*-995S) are at different nucleotide positions but affect the same codon and are mutually exclusive. We therefore re-coded these two SNPs as a single variant with three alleles: L (wild-type), F and S. Similarly, we also re-coded the two RDL SNPs as a single variant with three alleles.

Associations between genotype and insecticide resistance were tested using generalised linear models (GLM) with binomial errors and a logit link function, implemented in R using the package *lme4* (Bates et al. 2015). Genotypes were included as categorical fixed effects. Three candidate resistance markers (*Vgsc*-1853I, *Vgsc*-1934V and *Vgsc*-1874S) were not included in the models due to the scarcity of the mutant allele (*n* < 5) in the combined data, and the marker for *Vgsc*-791M was excluded as it was almost perfectly associated with *Vgsc*-1746S (Supplementary Table S1). For the data from Ghana, where all samples came from the same location, no random effects were included and the analysis was performed using the *glm* function. For the three other countries, sampling location was included as a random effect using generalised linear mixed models (GLMM) implemented by the function *glmer*. In one case (permethrin phenotype from DRC) where the mixed models failed to converge, preventing the inclusion of random effects, the modelling was repeated by including location as a fixed effect, using the function *glm*. In Burkina Faso and DRC, the majority of samples were of the species *An*. *gambiae, with* very few *An*. *coluzzii* (7 from Burkina Faso, 2 from the DRC), and *An*. *coluzzii* samples were thus excluded from the analyses. Similarly, only 6 *An*. *arabiensis* samples were found among the samples tested for resistance to deltamethrin in Kenya and were thus excluded. *An*. *arabiensis* were better represented in Kenya among the samples tested for resistance against permethrin (18 *An*. *arabiensis* samples), and species was thus included as a random factor.

Minimal significant models were obtained by a stepwise backward elimination process using a custom R function provided in the supplementary materials and described in Supplementary Methods S2. Briefly, resistance markers were included together into the full model, and the significance of each marker was obtained using the *anova* function to compare the full model to the model without the marker. Stepwise, the least significant marker was removed until only significant markers were left, providing the minimal model. The *P*-values reported for the significant markers are the result of the ANOVA comparing the minimal model against the model with the marker removed.

### 2.6 Detection of *Gste2* duplication

The Gstue_Dup7 duplication was detected by PCR using primers designed on either side of the duplication breakpoint. The position of the breakpoint was known from previous data (Lucas et al. 2018) and primers were designed using NCBI Primer Blast (Ye et al. 2012). Further details on the primers are provided in Supplementary Methods S3.

## 3. Results

### 3.1 High concordance between iPLEX MassARRAY and Ag1000G genotype calls

In total, 29,148 assays were run using the iPLEX MassARRAY panel (1,041 samples × 28 SNPs). Of these, 1,408 (4.8%) failed to produce a genotype call, with none of the individual assays having a failure rate greater than 10% (Supplementary Figure S1). The failure rate was lowest in *An*. *coluzzii*, where 24 out of 7,980 assays failed (0.3%, Supplementary Figure S2), followed by *An*. *gambiae*, where 274 out of 18,788 assays failed (1.46%, Supplementary Figure S3), and highest in *An*. *arabiensis*, where there were 59 failures out of 672 assays (8.8%, Supplementary Figure S4). The remaining samples were of unknown species as the DNA was not of high enough quality for the species ID assay.

Of the 8,036 individual assays performed on samples from Ag1000G (287 samples × 28 SNPs), only 5 (0.06%) iPLEX calls were divergent from the Ag1000G callset. The rest either gave concordant calls (98.2%) or failed calling by the iPLEX platform (1.7%). The high levels of concordance with the Ag1000G calls indicate that the results of the iPLEX assays can be used with confidence.

### 3.2 Deviation from Hardy-Weinberg equilibrium implies heterologous duplication of *Gste2* in Ghana

The SNPs *Ace-1* 280S and *Gste2* 114T showed significant deviation from Hardy-Weinberg expectations in *An*. *coluzzii* from Ghana, caused by an excess of heterozygotes at these loci (*P* < 10^−13^ in both cases, Table 1). Such heterozygote excess can be caused by the presence of heterologous amplifications, which can create functional heterozygotes by combining wild-type and mutant alleles on a single chromosome. For both *Gste2* 114T and *Ace-1* 280S, the raw intensities for heterozygous samples are split into several clusters rather than the normal single cluster (Fig. 1), which is also consistent with the presence of duplicate copies of one or both alleles. The heterologous *Ace-1* duplication in Ghana is already well-documented (Assogba et al. 2016, Weetman et al. 2015), but no such duplication has yet been reported in *Gste2*.

**Table 1:** Allele frequencies for *Gste2*-114T and Ace1-119S show large excesses of heterozygotes in Ghana. *Gste2*-114T: wild-type allele = A, mutant = G. *Ace1*-119S: wild-type allele = G, mutant = A.

|  |  | Burkina Faso<br>*An. gambiae* | DRC<br>*An. gambiae* | Ghana<br>*An. coluzzii* | Kenya<br>*An. gambiae* |
| --- | --- | --- | --- | --- | --- |
| *Gste2*-114T | AA | 71 | 148 | 3 | 139 |
|  | AG | 8 | 0 | 214 | 0 |
|  | GG | 1 | 0 | 21 | 0 |
| *Ace1*-280S | AA | 0 | 0 | 2 | 0 |
|  | AG | 33 | 0 | 153 | 0 |
|  | GG | 56 | 186 | 83 | 155 |

**Figure 1:**
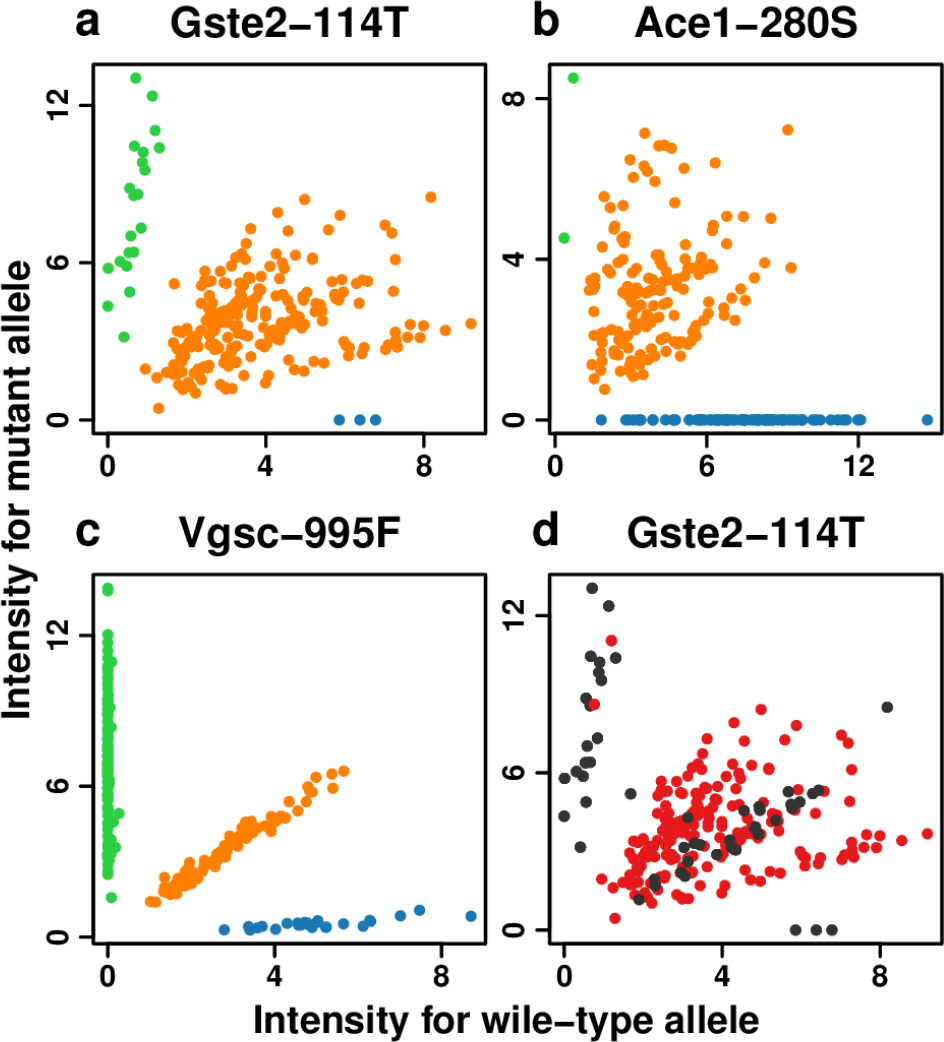
Raw intensities from the iPLEX MassARRAY assays in Ghana show multiple clusters among heterozygote samples (orange points) for *Gste2*-114T (a) and *Ace1*-280S (b). This is typically seen when duplications create ratios of mutant / wild-type alleles that differ from the usual 1/1 in heterozygotes. In comparison, equivalent data for *Vgsc*-995F (c) shows the normal pattern, with heterozygotes consistently falling in a single cluster along a straight diagonal line. Homozygote wild-type and homozygote mutant samples are shown in blue and red respectively Colour-coding the *Gste2*-114T data according to whether samples carry Dup7 (red) or not (black) shows that heterozygotes with no duplication predominantly fall along a straight diagonal line (d). The X and Y axes respectively show intensities for the wild-type and mutant alleles.

To confirm whether the heterozygote excess in *Gste2* 114T is caused by a duplication, we tested our samples for the presence of the duplication Gstue_Dup7, the only *Gste2* duplication known to occur in *An*. *coluzzii* from Ghana (Lucas et al. 2018). Gstue_Dup7 was present in 185 out of 238 (78%) of samples in our *An*. *coluzzii* samples from Ghana, a figure seven times higher than was found in Ag1000G, where 6 out of 55 (11%) *An*. *coluzzii* from Ghana had the duplication (Lucas et al. 2018).

Gstue_Dup7 was strongly association with heterozygosity at *Gste2* 114T (Fig. 1d). Out of 185 Ghanaian *An*. *coluzzii* samples that carried Gstue_Dup7, 183 were heterozygote at the 114T locus and 2 were homozygote mutants, while none were homozygote wild-type. In contrast, out of 53 samples that did not carry Gstue_Dup7, 31 were 114T heterozygotes, 19 were homozygote mutants and 3 were homozygote wild-type. Furthermore, the 114T heterozygotes without the duplication did not show the same scatter of raw intensities as the heterozygotes that carried the duplication (Fig. 1d).

### 3.3 High diversity of *kdr* haplotypes in the DRC

The *Vgsc*-995 mutations were at high frequency in all four countries (Fig. 2, Table 2). The wild-type allele was found at 2% frequency in *An*. *gambiae* from Kenya, but was completely absent in *An*. *gambiae* from Burkina Faso and the DRC. In *An*. *coluzzii*, the wild-type codon was found at 27% frequency in Ghana, and at similar frequencies in the few samples from Burkina Faso and the DRC.

**Table 2:**
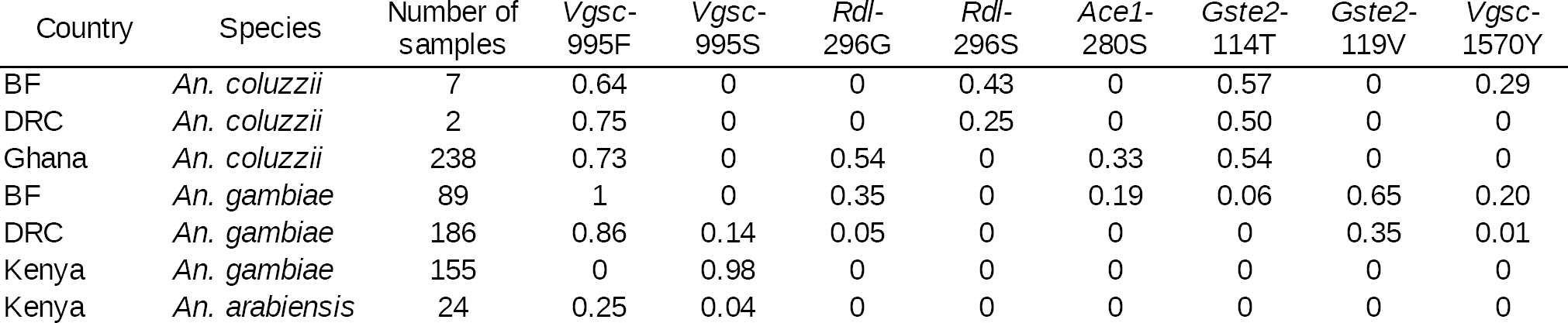
Allele frequencies for the eight SNPs with known associations with insecticide resistance. The full table showing all 28 markers in the panel is provided in Supplementary Data S5.

| Country | Species | Number of samples | Vgsc-995F | Vgsc-995S | Rdl-296G | Rdl-296S | Ace1-280S | Gste2-114T | Gste2-119V | Vgsc-1570Y |
| --- | --- | --- | --- | --- | --- | --- | --- | --- | --- | --- |
| BF | *An. coluzzii* | 7 | 0.64 | 0 | 0 | 0.43 | 0 | 0.57 | 0 | 0.29 |
| DRC | *An. coluzzii* | 2 | 0.75 | 0 | 0 | 0.25 | 0 | 0.50 | 0 | 0 |
| Ghana | *An. coluzzii* | 238 | 0.73 | 0 | 0.54 | 0 | 0.33 | 0.54 | 0 | 0 |
| BF | *An. gambiae* | 89 | 1 | 0 | 0.35 | 0 | 0.19 | 0.06 | 0.65 | 0.20 |
| DRC | *An. gambiae* | 186 | 0.86 | 0.14 | 0.05 | 0 | 0 | 0 | 0.35 | 0.01 |
| Kenya | *An. gambiae* | 155 | 0 | 0.98 | 0 | 0 | 0 | 0 | 0 | 0 |
| Kenya | *An. arabiensis* | 24 | 0.25 | 0.04 | 0 | 0 | 0 | 0 | 0 | 0 |

**Figure 2:**
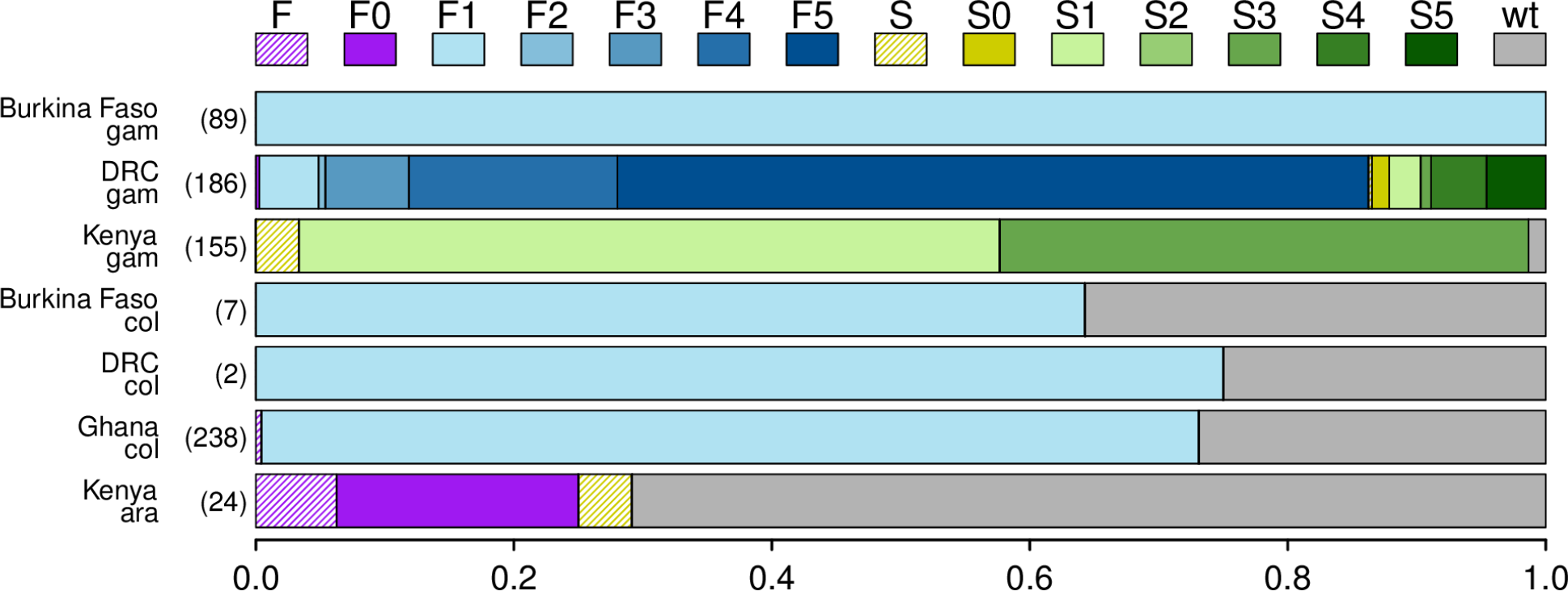
Frequencies of *kdr* haplotypes in the four countries from the study (total number of individuals, i.e. half the number of haplotypes, shown in brackets). The F1 haplotype is the only one present in Ghana and Burkina Faso, while haplotypes S1 and S3, along with a possible new *Vgsc*-995F haplotype, are found in Kenya. In the DRC, at least nine out of the ten *kdr* haplotypes co-exist. F and S indicate samples for which the *kdr* mutation was known, but the haplotype could not be determined because of failed calls in at least some background markers; F0 and S0 indicate haplotypes where the *kdr* mutation was known and all of the background markers produced wild-type calls, suggesting a novel haplotype background; wt = wild-type; ara = *An. arabiensis*; col = *An. coluzzii*; gam = *An. gambiae*.

In the West African populations (Burkina Faso and Ghana), all *kdr* mutant alleles were *Vgsc*- 995F of the F1 haplotype background (Fig. 2, Supplementary Data S6). In Kenya, all *kdr* mutants in *An*. *gambiae* carried the *Vgsc*-995S mutant allele from either of two backgrounds (S1 or S3; Supplementary Data S6). In contrast, most *kdr* haplotypes in Kenyan *An*. *arabiensis* were wild-type (34 out of 48), while the rest were predominantly *Vgsc*-995F (12 out of 48). These *Vgsc*-995F samples did not carry any of the SNPs associated with the five known haplotype background, possibly indicating that the *Vgsc*-995F haplotype background in *An*. *arabiensis* is distinct from those found to date in *An*. *gambiae*. The two *Vgsc*-995S haplotypes in this population could not be assigned a background due to failure of most of the haplotype background assays.

In the DRC, all five *Vgsc*-995F haplotype backgrounds were detected, the most frequent background being F5. Similarly, four of the five *Vgsc*-995S backgrounds were found in the DRC (Fig. 2, Supplementary Data S6), the most frequent background being S5, and haplotype S2 being absent. Previously, eight *kdr* haplotypes were found in Cameroon, with haplotypes S1 and S3 being absent (Anopheles gambiae 1000 Genomes Consortium 2017).

### 3.4 Association with insecticide resistance

In Burkina Faso, both *Gste2* mutations were associated with increased resistance to permethrin, demonstrating for the first time an association between *Gste2* 119V and insecticide resistance (Table 3). In contrast, the 1874L *Vgsc* mutation was negatively associated with resistance to permethrin.

**Table 3:** Significant associations between genotype and insecticide resistance. Effects are shown as log odds ratio. Direction of effect compared to the reference (intercept) genotype shown by arrows (↑ indicates increase in resistance compared to the intercept; ↓ indicates decrease in resistance). In Ghana, *kdr* was either wild-type (L) or 995F. In the DRC, *kdr* was either 995S or 995F.

| Country | Insecticide | Codon | P | Intercept genotype | Effect of het. vs intercept | Effect of hom. mutant vs intercept | Effect of hom. mutant vs het. |
| --- | --- | --- | --- | --- | --- | --- | --- |
| Ghana | Perm | Vgsc-995 | 0.023 | LL | LF: ≥ 4* ↑ | FF: ≥ 4* ↑ | 1.58 ↑ |
|  | Delt | NA |  |  |  |  |  |
|  | DDT | Rdl-296 | 7.1x10<sup>-7</sup> | AA | AG: 2.6 ↑ | GG: 2.95 ↑ | 0.35 ↑ |
| Burkina Faso | Perm | Vgsc-1874L | 0.041 | PP | LP: -1.18 ↓ | LL: -2.4 ↓ | -1.22 ↓ |
|  |  | Gste2-114T | 0.022 | II | IT: 3.01 ↑ | TT: ≥ 4* ↑ | ≥ 4* ↑ |
|  |  | Gste2-119V | 0.026 | LL | LV: ≥ 4* ↑ | VV: ≥ 4* ↑ | 0.35 ↑ |
| Kenya | Perm | NA |  |  |  |  |  |
|  | Delt | NA |  |  |  |  |  |
| DRC | Perm | Vgsc-995 | 0.004 | SS | SF: ≥ 4* ↑ | FF: ≥ 4* ↑ | 1.1 ↑ |
|  | Delt | Gste2-119V | 0.0078 | LL | LV: 0.36 ↑ | VV: -1.54 ↓ | -1.91 ↓ |
|  |  | Vgsc-995 | 0.0038 | SS | SF: -0.55 ↓ | FF: 1.5 ↑ | 2.06 ↑ |

\* Effects ≥ 4 (corresponding to odds ratio > 50) were usually seen where there were few samples in a category, leading to all samples having the same phenotype and making odds ratios impossible to calculate.

In the DRC, where the wild-type *Vgsc*-995 codon was absent, the *Vgsc*-995F mutation was more strongly associated with resistance to both permethrin and deltamethrin than *Vgsc*-995S, with a slight tendency for heterozygotes to have lower resistance to deltamethrin than *Vgsc*-995S homozygotes (Table 3). Seemingly contrary to the Burkina Faso data, there was a negative association between the *Gste2*-119V mutation and resistance to deltamethrin, although heterozygotes showed slightly higher resistance than homozygote wild-types.

In Ghana, no SNPs were associated with resistance to deltamethrin, while the *Vgsc*-995F mutation was associated with resistance to permethrin and the *Rdl*-296G mutation was associated with resistance to DDT (Table 3). There were only three homozygous wild-type samples at the *Vgsc*-995F locus, thus the significant difference is driven by the difference between the homozygote mutant and heterozygotes.

In Kenya, there were no significant associations between any of the SNPs and resistance to either permethrin or deltamethrin, although it is notable that *Vgsc*-995 was the only resistance candidate SNP found in this population and was nearly fixed in *An*. *gambiae* (Table 2 & Supplementary Data S5).

## 4 Discussion

We have developed a multiplex SNP panel for quick, high-throughput screening of large numbers of *An*. *gambiae s*.*l*. at multiple markers useful for detecting insecticide resistance, testing the importance of putative resistance diagnostic markers and tracking the spread of resistance mutations. We have allied this with a formal analytical approach for analysing phenotypic association of these markers, made available as an R function. By screening 713 mosquitoes from four different countries, we have identified a heterologous duplication in a resistance-conferring SNP, confirmed an association with resistance in a candidate SNP, and found the DRC to be an area in which nearly all evolutionary origins of the *kdr* resistance mutation co-exist.

The presence of a heterologous duplication in *Gste2* in *An*. *gambiae* from Ghana was first indicated by a very high excess in *Gste2*-114T heterozygotes and the characteristic multiple clusters of signal intensities in the raw data, both of which bear a striking resemblance to the pattern seen in Ghanaian *Ace1*-280S data, known to be caused by a heterologous duplication. We confirmed the role of a duplication by showing that the heterozygosity is associated with the presence of the *Gste2* duplication Gstue_Dup7 (Lucas et al. 2018). While the vast majority of Gstue_Dup7 alleles are heterologous, we also found evidence of a rare variant of Gstue_Dup7 in two samples where the duplication was homozygous for the 114T SNP. So far, eleven different duplications in the cluster of genes around *Gste2* have been reported in *An*. *gambiae* and *An*. *coluzzii* from across SSA (Lucas et al. 2018), but only one of these, Gstue_Dup1, was shown to be associated with the 114T mutation, being homologous for the mutant allele. The evolution of duplications in *Gste2* may therefore mirror what has been found in *Ace-1*, where both homologous and heterologous duplications exist. If the 114T mutation carries a physiological cost in the absence of insecticide pressure, the heterologous duplication discovered here may serve to reduce this cost, as has been found in the case of the *Ace-1* duplication (Assogba et al. 2015).

Gstue_Dup7 was found in 78% of our *An*. *coluzzii* samples from Ghana collected in 2015 from Keta (5.9175N, 0.9916E), which is seven times higher than was found in samples collected in 2012, albeit from different sites further West in Ghana (Koforidua (6.0945N, 0.2609W), Madina (5.6685N, 0.2193W), Takoradi (4.9122N, 1.7740W), Twifo Praso (5.6086N, 1.5493W); Lucas et al. 2018). While we cannot exclude the possibility that this increase reflects differences between sites, the high frequency of this duplication in Keta, combined with the striking difference in frequency between samples collected in 2012 and 2015, strongly suggest that this duplication is rapidly spreading through the *An*. *coluzzii* population.

The high diversity of *kdr* haplotype backgrounds in the DRC points to this area of central Africa as a key to understanding the spread of the most well-characterised insecticide resistance allele in *An*. *gambiae*. All five *Vgsc*-995F backgrounds were found in this population (*Anopheles gambiae* 1000 Genomes Consortium 2017), as well as four of the five *Vgsc*-995S backgrounds, combining haplotypes previously found in Uganda, (S1 and S3), Kenya (S3) and Cameroon (S4 and S5). The only background not confirmed in the DRC was S2, which was previously found in Gabon and Cameroon (*Anopheles gambiae* 1000 Genomes Consortium 2017). This striking geographical co-existence of *kdr* haplotype backgrounds could have two explanations. First, most of the main *kdr* origins could have occured in central Africa, and subsequently spread from there to other countries, with different haplotypes becoming established in different regions. Second, The different *kdr* haplotypes could have spread from their respective origins and converged in the DRC, where most then persisted without being entirely replaced (although F5 is by far the most prevalent, with a frequency of over 50%). Differentiating between these two hypotheses will require more in-depth analysis of the sequence variability within each haplotype sequence for each background in the DRC and elsewhere.

The *Vgsc*-995F haplotype found in *An*. *arabiensis* from Kenya did not conform to any of the known haplotype backgrounds for this mutation, suggesting that a sixth *Vgsc*-995F origin exists, which may have originated in East Africa. The Kenyan samples in the present study are from Western Kenya, where *Vgsc*-995F has previoulsy been reported in both *An*. *arabiensis* and *A*. *gambiae* (Ochomo et al. 2015). In contrast, the Kenyan samples included in the Ag1000G data, where the haplotype backgrounds were defined, came from an eastern Kenyan population of *An*. *gambiae* where *kdr*-995F was not found. The Kenyan *Vgsc*-995F background was therefore not present in Ag1000G and could not be included in the definition of haplotype backgrounds. Whether the *Vgsc*-995F background present in *An*. *gambiae* from western Kenya corresponds to one of the five known 995F backgrounds, or to the one found in *An*. *arabiensis* from the same area remains to be resolved.

Associations of genetic markers with insecticide resistance revealed several unexpected associations. The *Gste2*-114T marker, which has been previously associated with resistance to DDT (Mitchell et al. 2014) was associated with resistance to permethrin in Burkina Faso. To our knowledge, this is the first report of a role for this mutation in permethrin resistance, although one previous study found that it was also associated with resistance to another pyrethroid, deltamethrin (Opondo et al. 2016). The evidence is therefore increasing for a role of *Gste2*-114T in cross-resistance to multiple insecticides. In Ghana, there was no significant association between *Gste2*-114T and resistance to pyrethroids or DDT, but this is likely due to the very high frequency of heterozygotes, making any association difficult to detect. In the same gene, the *Gste2*-119V mutation was associated with increased permethrin resistance in Burkina Faso, but with decreased deltamethrin resistance in the DRC. These contrasting effects may reflect different changes in binding affinity to these two insecticides caused by this mutation. While the metabolic properties induced by the *Gste2*-119V mutation have not been studied, research on the *Gste2*-119F mutation in *An*. *funestus* showed that the mutant form was able to metabolise permethrin, but not deltamethrin (Riveron et al. 2014). The negative association with deltamethrin resistance found here is, however, novel, and may be explained by a metabolic cost imposed by the mutation, or by reduced affinity for deltamethrin in the case of the *An*. *gambiae* mutation. If the opposing effects on permethrin and deltamethrin resistance detected here are confirmed, it will be important to routinely screen for this mutation when making a choice of insecticides in control campaigns.

Intriguingly, *Rdl*-296G was associated with resistance to DDT in Ghana. This mutation is in the target site of cyclodiene insecticides such as dieldrin, which was used in malaria control programmes in Africa during the 1950s and 1960s, but has since been discontinued. Resistance to dieldrin has somehow been maintained in natural mosquito populations (Asih et al. 2012, Kwiatkowska et al. 2013, Wondji et al. 2011), although the selective forces causing its persistence remain a puzzle. One possibility is that cyclodienes continue to be used as insecticides in private agriculture, but another is that the *Rdl*-296 mutations confer cross-resistance to other insecticides. For example, insertion of the *An*. *gambiae Rdl* gene into *Xenopus* oocytes demonstrated that fipronil, deltamethrin and imidacloprid all inhibited the GABA receptor, of which RDL is a subunit, and that this effect was reduced by the *Rdl*-296G mutation (Taylor-Wells et al. 2015). However, the same experiment found no effect of DDT on RDL. Our results may point to an as-yet unrecognised example of cross-resistance between dieldrin and DDT, through mechanisms that remain to be elucidated.

The *Vgsc*-1874L mutation, exclusively found on the background of the *kdr*-995 F1 haplotype, was negatively associated with resistance to permethrin in Burkina Faso, where the F1 haplotype is fixed in the population. The mutation’s continued presence in the population despite its cost to permethrin resistance might be due to one of two factors. First, the mutation may provide improved resistance to other insecticides. However, since *kdr* is a target site resistance mechanism, the insecticides that this could apply to are limited to the pyrethroids and DDT. Second, the *Vgsc*-1874L mutation may serve to compensate the costs of *kdr*, serving to increase the fitness of the haplotype in the absence of insecticides. This would be a worrying development in the evolution of insecticide resistance, as it would favour the persistence of the *kdr* mutation during cycles of insecticide rotation in control programmes.

In conclusion, our results highlight the importance and value of high-throughput genetic screening for the study and monitoring of insecticide resistance in malaria vectors. The three main steps involved in developing new markers of insecticide resistance are 1. discovery of new mutations, 2. confirmation of association with resistance and 3. tracking the mutation across populations. Our panel of 28 genetic markers achieved all three of these steps, albeit on different markers. We found evidence for a heterologous duplication in *Gste2*, we reported the first evidence of resistance association for *Gste2*-119V, bringing the total number of resistance-associated SNPs in *An*. *gambiae s*.*l*. up to 10, and we investigated the distribution of *Vgsc*-995 mutations across East, Central and West Africa. This work should now expand in two directions. First, the panel should be applied to more populations, building a more complete picture of the distribution of these genotypes and their importance in resistance. Second, more markers should be gradually added to the panel as their importance becomes recognised. Once routine monitoring of important mutations become common-place, our understanding of the genetic underpinnings of insecticide resistance will increase dramatically.

## Supporting information

Supplementary R functions

## 5 Acknowledgements

Funding was provided by Award Number R01AI116811 from the National Institute of Allergy and Infectious Diseases (NIAID) and Award MR/P02520X/1 from the Medical Research Council, UK. The work was also supported by the EC FP7 Project grant no: 265660 “AvecNet” and a Wellcome Trust also core award to The Wellcome Trust Centre for Human Genetics (090532/Z/09/Z, 203141). KR and CH are supported by the Wellcome Trust (090770/Z/09/Z, 204911/Z/16/Z).

