## Supplementary Figures and Tables for "A high throughput multi-locus insecticide resistance marker panel for tracking resistance emergence and spread in *Anopheles gambiae*"

### Electronic Supplementary Material
### Supplementary figures

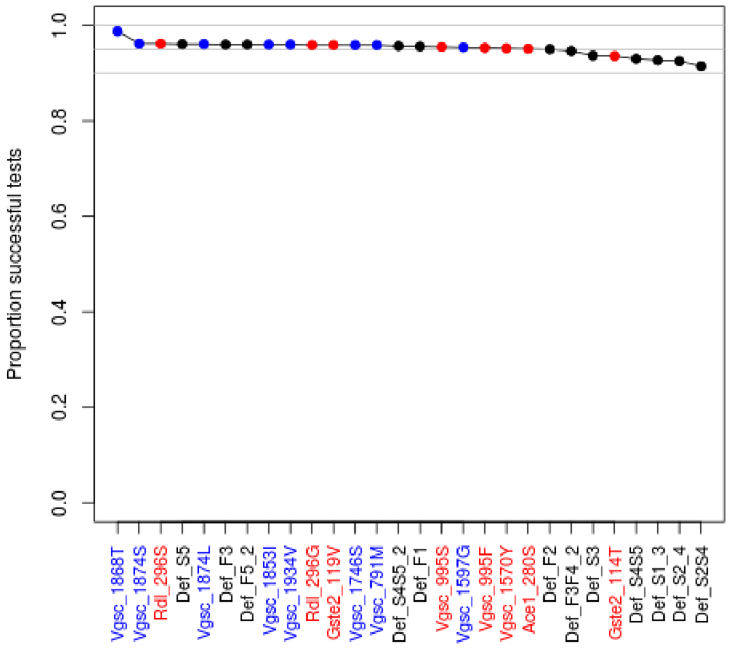


**Fig. S1**: Success rate of individual SNP assays in the multiplex sequenome assay was never lower than 90% (measured as the proportion of assays that were of high enough quality to give a genotype call). Red points indicate known resistance variants, blue points indicate variants found on the background of some *kdr* mutations and which may therefore be associated with either increased resistance or compensation for the costs of the *kdr* mutations, black points indicate SNPs used to distinguish between *kdr* mutant haplotype backgrounds.

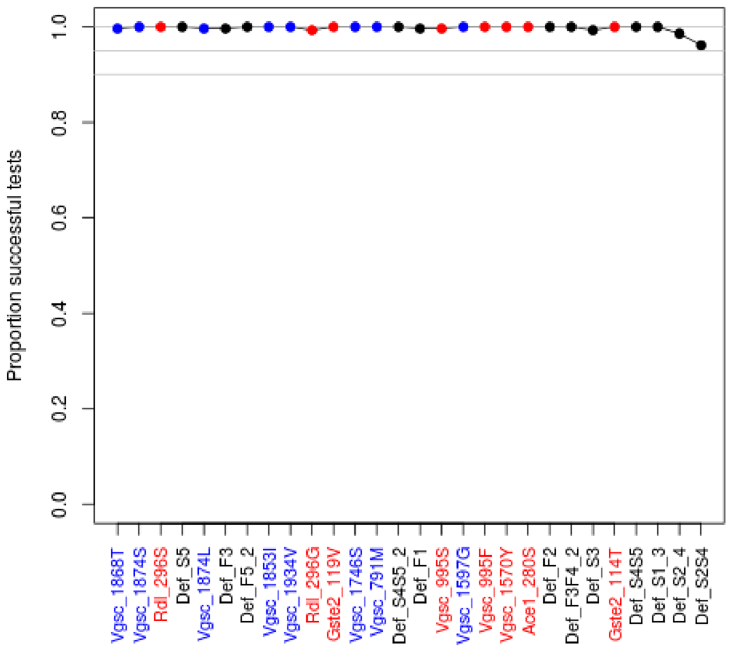


**Fig. S2**: Success rate of individual SNP assays among *An. coluzzii* samples was never lower than 95% (measured as the proportion of assays that were of high enough quality to give a genotype call). Red points indicate known resistance variants, blue points indicate variants found on the background of some *kdr* mutations, black points indicate SNPs used to distinguish between *kdr* mutant haplotype backgrounds.

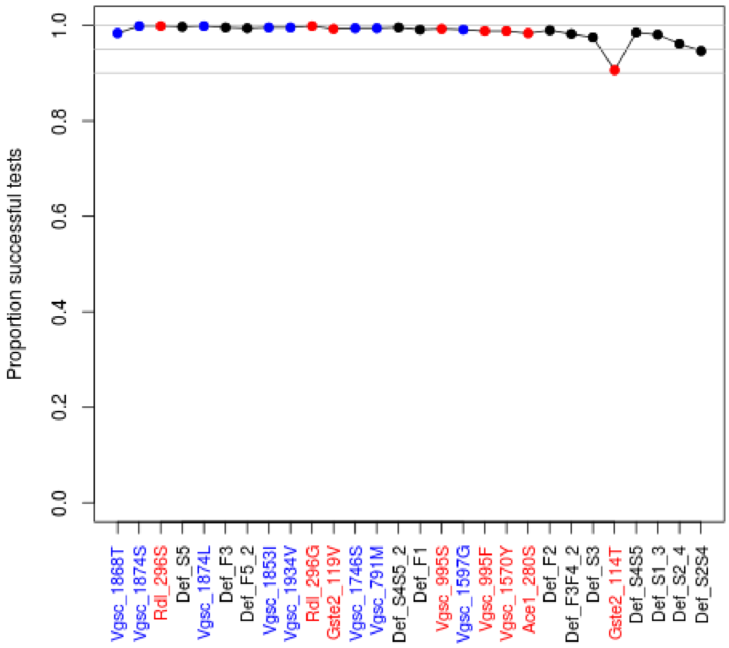


**Fig. S3**: Success rate of individual SNP assays among *An. gambiae* samples was never lower than 90% (measured as the proportion of assays that were of high enough quality to give a genotype call). Red points indicate known resistance variants, blue points indicate variants found on the background of some *kdr* mutations, black points indicate SNPs used to distinguish between *kdr* mutant haplotype backgrounds.

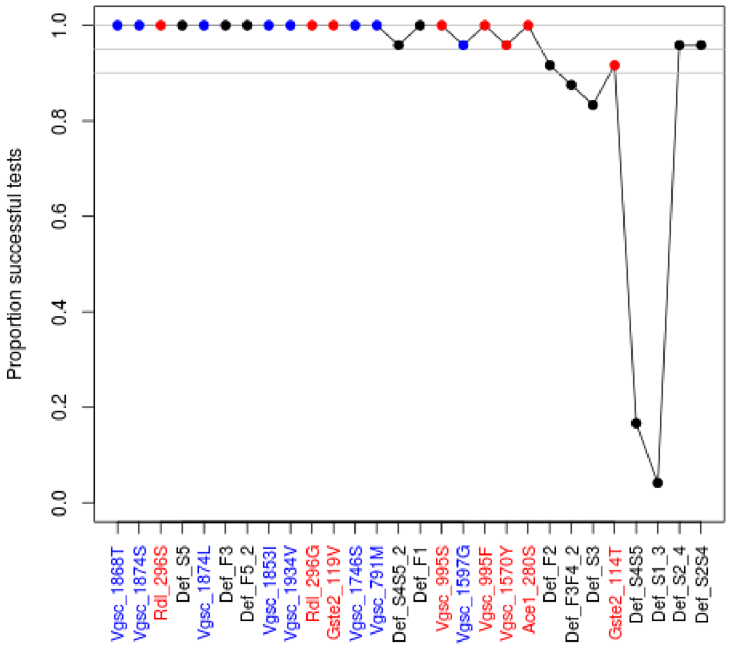


**Fig. S4**: Success rate of individual SNP assays among *An. arabiensis* samples was good overall, but some assays had a low rate of success (measured as the proportion of assays that were of high enough quality to give a genotype call). All four assays with success rates below 90% were ones used to distinguish between *kdr* haplotype backgrounds Red points indicate known resistance variants, blue points indicate variants found on the background of some *kdr* mutations, black points indicate SNPs used to distinguish between *kdr* mutant haplotype backgrounds.

Table **S1**: SNPs VGC-1746S and VGC-791M are almost perfectly associated (only two samples give discordant calls). The wild-type alleles are G and C for 1746S and 791M respectively.

| | | VGC-1746S | | |
|---|---|---|---|---|
| | | GG | GT | TT |
| | CC | 672 | 0 | 0 |
| VGC-791M | CT | 2 | 33 | 0 |
| | TT | 0 | 0 | 5 |
