## Supplementary Methods for "A high throughput multi-locus insecticide resistance marker panel for tracking resistance emergence and spread in *Anopheles gambiae*"

### Electronic Supplementary Material

### Supplementary methods

### S1 Implementation of kdr haplotype background markers.

Background markers were only taken into consideration if the associatied *kdr* mutation was already detected using the two kdr markers (Vgsc_995F and Vgsc_995S). Where a sample was found to carry the *Vgsc*-995F allele, the five 995F haplotype background markers (Def_F1, Def_F2, Def_F3F4_2, Def_F3, Def_F5_2) were used to determine that allele's haplotype background. Markers Def_F1, Def_F2 and Def_F5_2 were applied using the method shown in Fig. **M1a**. Because no marker could be found that defined haplotype F4, we instead used marker Def_F3F4 to separate the F3 and F4 backgrounds from the other 995F backgrounds. Samples positive for the (F3F4) background were then separated into F3 and F4 using marker Def_F3 (Fig. **M1b**).

Where a sample was found to carry the kdr995S allele, the seven 995S haplotype background markers (Def_S1_3, Def_S2_4, Def_S2S4, Def_S3, Def_S5 Def_S4S5 and Def_S4S5_2) were used to determine that allele's haplotype background. Markers Def_S1_3, and Def_S3 were applied using the method shown in Fig. **M1a**. Because no marker could be found that defined haplotype S4, we instead used pairs of markers as described above for haplotype background F4 (Fig. **M1b**). We found one marker that could separate the S2 and S4 backgrounds from the other 995S backgrounds (Def_S2S4), which could be combined with Def_S2_4 to identify S2 and S4. We also found two markers that could separate the S4 and S5 backgrounds from the other 995S backgrounds (Def_S4S5 and Def_S4S5_2), which could be combined with Def_S5 to identify S2 and S5, thereby providing independent confirmation of a 995S allele belonging to the S4 haplotype background.).

Where a sample was reported to be homozygous for one of the *kdr* haplotype backgrounds, but only heterozygous for the kdr allele itself, the process was considered to have failed and no haplotype background was reported for the sample.

#### *Vgsc*-995F background calls in the DRC

In some cases, while a marker distinguishes one haplotype background from the others with the same *kdr* mutation, that same marker is also variable on non-*kdr* backgrounds, or on the background of the other *kdr* mutant (Supplementary Data S2). For example, marker Def_F2 distinguishes the *kdr* F2 background from the other F backgrounds, but the F2 allele is also found at 55% frequency on the S background and at 17% frequency on the wild-type background in the Ag1000G data. Therefore, samples that are heterozygous for 995F can appear to have two different haplotype backgrounds if the non-995F haplotype carries the F2 allele at the Def_F2 marker. Such samples were found in our data from the DRC, where 995F heterozygotes were called as both F2 and one of F3, F4 or F5. The nine *Vgsc*-995 homozygote wild-type samples in the DRC never carried SNPs associated with F3 or F4, and only two samples were heterozygote for the F5 SNP (equating to an allele frequency of 11%). In contrast, the frequency of the F2 SNP in these samples was 67%. Furthermore, among *Vgsc*-995F homozygotes, the F5 background was by far the most frequent in the DRC (Supplementary Data S6). We therefore considered that, where the *Vgsc*-995F heterozygotes appeared to have both the F2 and another haplotype background, the F2 background was incorrect.

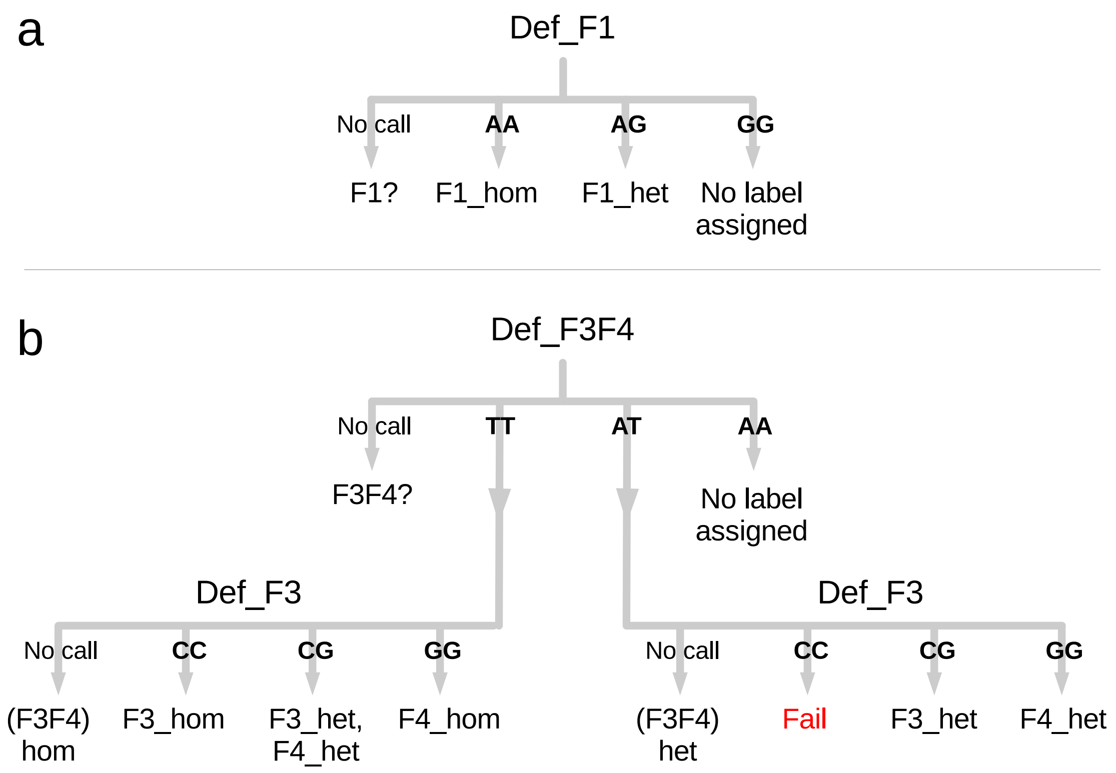


**Fig. M1**: Process used to interpret the kdr haplotype background markers for each sample. This process was implemented using the custom R functions *F.kdr.origin* and *S.kdr.origin* provided in the file `Supplementary_Materials_R_functions.r`. *het* = heterozygous; *hom* = homozygous. **a** Markers that defined a single kdr haplotype background were assessed using a simple decision tree such as the one depicted here for the marker defining the F1 haplotype background (Def_F1). A genotype call of AT and TT at this marker result in the sample being labelled as heterozygous and homozygous respectively for the F1 background. A genotype call of AA results in no label being applied to the sample. **b** Pairs of markers that acted together to defined kdr haplotype backgrounds were assessed using a decision tree such as the one depicted here for the markers Def_F3F4 and Def_F3. In this example, the marker Def_F3F4 identifies the F3 and F4 backgrounds from among the 995F backgrounds. A seperate marker, Def_F3, identifies F3 samples, thus differentiating F3 and F4. Contradictory situations, such as where the sample is homozygous for the Def_F3 markers but only heterozygous for the Def_F3F4 marker, result in the sample failing the haplotype background calling process.

### S2 Statistical analysis of association between SNPs and insecticide resistance.

The custom R function used to perform this analysis (*glmodelling*) has been provided in the file `Supplementary_Materials_R_functions.r`.

The process of the function is verbally summarised as follows::

1. Verify that any random effect terms provided to the function have more than one value in the data. Remove any that do not. All models used in the rest of the function, including the null model, will include the remaining random effect terms.

2. Verify that any SNP markers provided to the function are sufficiently variable (ie: at least two samples do not have the most frequent genotype). Remove any that are not, and report an error if no sufficiently variable markers are left.

3. Use a generalised linear model to investigate each marker individually by comparing the model with the marker removed to the null model. Store these results separately, they will not be used in the model reduction process.

4. Enter a loop that will end once the minimal model is reached:

   (a) Where at least one marker is NA (has no genotype call) in a sample, the glm will ignore that sample completely. In doing so, it is possible that other markers then become monomorphic once that sample is removed, which will cause the glm model to fail. Therefore, we remove markers that would become monomorphic, but only for the duration of this iteration of the loop.

   (b) Build the glm model including all of the remaining markers.

   (c) For each marker, build a reduced model with that marker removed. Use the *anova* function to compare it to the model in 4b and obtain a $P$-value for the removed marker. If the reduced model contains more samples than the model in 4b, then it needs to be built with those samples removed to allow the comparison. If the model gives no $P$-value, this indicates that the marker is perfectly correlated with another marker. We therefore immediately remove this marker and start a new iteration of the loop (ie: return to 4a).

   (d) If all markers have a $P$-value below 0.05, accept this as the minimal model. End the loop and provide this model as the output to the function.

   (e) Otherwise, remove the marker with the highest $P$-value and start a new iteration of the loop (ie: return to 4a) or, if no markers remain, report the null model as the minimal model. End the loop and provide the null model as the output to the function.

### S3 PCR for detection of Gstue_Dup7.

PCR primers (Table **M1**) were designed either side of the Gstue_Dup7 breakpoint using the breakpoint sequence reported in Lucas et al. (2018), such that the primers would create a product only in the presence of the duplication. To differentiate between the absence of a duplication and PCR failure, a third primer was also added to the reaction that amplifies with the forward primer even in the absence of the duplication.

Table **M1**: Primer sequences used to detect Gstue_Dup7. Primers Ag_Gst_Dup7_1F and Ag_Gst_Dup7_1R combine to create a 151bp product when the duplication is present. Primers Ag_Gst_Dup7_1F and Ag_Gst_Dup7_1Rc2 combine to create a 265bp product regardless of the presence of the duplication, providing a test that the PCR was successful.

| Primer name | Primer sequence |
|---|---|
| Ag_Gst_Dup7_1F | TCGAACGAACCCCACGATTT |
| Ag_Gst_Dup7_1R | GCGGCCCTCTGATGAAATGA |
| Ag_Gst_Dup7_1Rc2 | CCCGAACGCGTAACGTAAAC |
