## Supplementary R functions for "A high throughput multi-locus insecticide resistance marker panel for tracking resistance emergence and spread in *Anopheles gambiae*"

library(modEvA)
library(lme4)

# For a given sample, determine its kdr F origin.
F.kdr.origin <- function(sample.row){
	# Check for the Vgsc-995F mutation (recorded as Ag_kdr_1014F in the table)
	if (is.na(sample.row[,'Ag_kdr_1014F']))
		kdr.F.origins <- 'F:?'
	else if (sample.row[,'Ag_kdr_1014F'] == 'AA')
		kdr.F.origins <- 'F:wt_hom'
	else if (sample.row[,'Ag_kdr_1014F'] == 'AT')
		kdr.F.origins <- 'F:het'
	else if (sample.row[,'Ag_kdr_1014F'] == 'TT')
		kdr.F.origins <- 'F:hom'
	# If the individual has the Vgsc-995F mutation, find out its haplotype background
	# for F homozygotes
	if (kdr.F.origins == 'F:hom'){
		if (is.na(sample.row[,'Ag_Def_F1']))
			kdr.F.origins <- paste(kdr.F.origins, 'F1?', sep = ',')
		else if (sample.row[,'Ag_Def_F1'] == 'AA')
			kdr.F.origins <- paste(kdr.F.origins, 'F1_hom', sep = ',')
		else if (sample.row[,'Ag_Def_F1'] == 'AG')
			kdr.F.origins <- paste(kdr.F.origins, 'F1_het', sep = ',')
		#
		if (is.na(sample.row[,'Ag_Def_F2']))
			kdr.F.origins <- paste(kdr.F.origins, 'F2?', sep = ',')
		else if (sample.row[,'Ag_Def_F2'] == 'AA')
			kdr.F.origins <- paste(kdr.F.origins, 'F2_hom', sep = ',')
		else if (sample.row[,'Ag_Def_F2'] == 'AG')
			kdr.F.origins <- paste(kdr.F.origins, 'F2_het', sep = ',')
		#
		if (is.na(sample.row[,'Ag_Def_F3F4_2']))
			kdr.F.origins <- paste(kdr.F.origins, 'F3F4?', sep = ',')
		else if (sample.row[,'Ag_Def_F3F4_2'] == 'TT'){
			if (is.na(sample.row[,'Ag_Def_F3']))
				kdr.F.origins <- paste(kdr.F.origins, '(F3F4)_hom', sep = ',')
			else if (sample.row[,'Ag_Def_F3'] == 'CC')
				kdr.F.origins <- paste(kdr.F.origins, 'F3_hom', sep = ',')
			else if (sample.row[,'Ag_Def_F3'] == 'CG')
				kdr.F.origins <- paste(kdr.F.origins, 'F3_het,F4_het', sep = ',')
			else if (sample.row[,'Ag_Def_F3'] == 'GG')
				kdr.F.origins <- paste(kdr.F.origins, 'F4_hom', sep = ',')
		}
		else if (sample.row[,'Ag_Def_F3F4_2'] == 'AT'){
			if (is.na(sample.row[,'Ag_Def_F3']))
				kdr.F.origins <- paste(kdr.F.origins, '(F3F4)_het', sep = ',')
			else if (sample.row[,'Ag_Def_F3'] == 'CC')
				return(paste('Fail. Genotypes suggest that sample', rownames(sample.row), 'is heterozygote for F3F4, but homozygote for F3.'))
			else if (sample.row[,'Ag_Def_F3'] == 'CG')
				kdr.F.origins <- paste(kdr.F.origins, 'F3_het', sep = ',')
			else if (sample.row[,'Ag_Def_F3'] == 'GG')
				kdr.F.origins <- paste(kdr.F.origins, 'F4_het', sep = ',')
		}
		#
		if (is.na(sample.row[,'Ag_Def_F5_2']))
			kdr.F.origins <- paste(kdr.F.origins, 'F5?', sep = ',')
		else if (sample.row[,'Ag_Def_F5_2'] == 'GG')
			kdr.F.origins <- paste(kdr.F.origins, 'F5_hom', sep = ',')
		else if (sample.row[,'Ag_Def_F5_2'] == 'AG')
			kdr.F.origins <- paste(kdr.F.origins, 'F5_het', sep = ',')
	}
	# for F heterozygotes
	else if (kdr.F.origins == 'F:het'){
		if (is.na(sample.row[,'Ag_Def_F1']))
			kdr.F.origins <- paste(kdr.F.origins, 'F1?', sep = ',')
		else if (sample.row[,'Ag_Def_F1'] == 'AA')
			return(paste('Fail. Genotypes suggest that sample', rownames(sample.row), 'is heterozygote for F kdr, but homozygote for F1.'))
		else if (sample.row[,'Ag_Def_F1'] == 'AG')
			kdr.F.origins <- paste(kdr.F.origins, 'F1_het', sep = ',')
		#
		if (is.na(sample.row[,'Ag_Def_F2']))
			kdr.F.origins <- paste(kdr.F.origins, 'F2?', sep = ',')
		else if (sample.row[,'Ag_Def_F2'] == 'AA')
			return(paste('Fail. Genotypes suggest that sample', rownames(sample.row), 'is heterozygote for F kdr, but homozygote for F2.'))
		else if (sample.row[,'Ag_Def_F2'] == 'AG')
			kdr.F.origins <- paste(kdr.F.origins, 'F2_het', sep = ',')
		#
		if (is.na(sample.row[,'Ag_Def_F3F4_2']))
			kdr.F.origins <- paste(kdr.F.origins, 'F3F4?', sep = ',')
		else if (sample.row[,'Ag_Def_F3F4_2'] == 'TT')
			return(paste('Fail. Genotypes suggest that sample', rownames(sample.row), 'is heterozygote for F kdr, but homozygote for F3F4.'))
		else if (sample.row[,'Ag_Def_F3F4_2'] == 'AT'){
			if (is.na(sample.row[,'Ag_Def_F3']))
				kdr.F.origins <- paste(kdr.F.origins, '(F3F4)_het', sep = ',')
			else if (sample.row[,'Ag_Def_F3'] == 'CC')
				return(paste('Fail. Genotypes suggest that sample', rownames(sample.row), 'is heterozygote for F kdr and F3F4, but homozygote for F3.'))
			else if (sample.row[,'Ag_Def_F3'] == 'CG')
				kdr.F.origins <- paste(kdr.F.origins, 'F3_het', sep = ',')
			else if (sample.row[,'Ag_Def_F3'] == 'GG')
				kdr.F.origins <- paste(kdr.F.origins, 'F4_het', sep = ',')
		}
		#
		if (is.na(sample.row[,'Ag_Def_F5_2']))
			kdr.F.origins <- paste(kdr.F.origins, 'F5?', sep = ',')
		else if (sample.row[,'Ag_Def_F5_2'] == 'GG')
			return(paste('Fail. Genotypes suggest that sample', rownames(sample.row), 'is heterozygote for F kdr, but homozygote for F5.'))
		else if (sample.row[,'Ag_Def_F5_2'] == 'AG')
			kdr.F.origins <- paste(kdr.F.origins, 'F5_het', sep = ',')
	}
	kdr.F.origins
}

# For a given sample, determine its kdr S origin. 
S.kdr.origin <- function(sample.row, alternate_S4S5 = F){
	# Check for the Vgsc-995S mutation (recorded as Ag_kdr_1014S in the table)
	if (is.na(sample.row[,'Ag_kdr_1014S']))
		kdr.S.origins <- 'S:?'
	else if (sample.row[,'Ag_kdr_1014S'] == 'TT')
		kdr.S.origins <- 'S:wt_hom'
	else if (sample.row[,'Ag_kdr_1014S'] == 'CT')
		kdr.S.origins <- 'S:het'
	else if (sample.row[,'Ag_kdr_1014S'] == 'CC')
		kdr.S.origins <- 'S:hom'
	# If the individual has kdr S, find out its origins
	# for S homozygotes
	if (kdr.S.origins == 'S:hom'){
		if (is.na(sample.row[,'Ag_Def_S1_3']))
			kdr.S.origins <- paste(kdr.S.origins, 'S1?', sep = ',')
		else if (sample.row[,'Ag_Def_S1_3'] == 'CC')
			kdr.S.origins <- paste(kdr.S.origins, 'S1_hom', sep = ',')
		else if (sample.row[,'Ag_Def_S1_3'] == 'CT')
			kdr.S.origins <- paste(kdr.S.origins, 'S1_het', sep = ',')
		#
		if (is.na(sample.row[,'Ag_Def_S2S4']))
			kdr.S.origins <- paste(kdr.S.origins, 'S2S4?', sep = ',')
		else if (sample.row[,'Ag_Def_S2S4'] == 'TT'){
			if (is.na(sample.row[,'Ag_Def_S2_4']))
				kdr.S.origins <- paste(kdr.S.origins, '(S2S4)_hom', sep = ',')
			else if (sample.row[,'Ag_Def_S2_4'] == 'AA')
				kdr.S.origins <- paste(kdr.S.origins, 'S2_hom', sep = ',')
			else if (sample.row[,'Ag_Def_S2_4'] == 'AT')
				kdr.S.origins <- paste(kdr.S.origins, 'S2_het,S4_het', sep = ',')
			else if (sample.row[,'Ag_Def_S2_4'] == 'TT')
				kdr.S.origins <- paste(kdr.S.origins, 'S4_hom', sep = ',')
		}
		else if (sample.row[,'Ag_Def_S2S4'] == 'CT'){
			if (is.na(sample.row[,'Ag_Def_S2_4']))
				kdr.S.origins <- paste(kdr.S.origins, '(S2S4)_het', sep = ',')
			else if (sample.row[,'Ag_Def_S2_4'] == 'AA')
				return(paste('Fail. Genotypes suggest that sample', rownames(sample.row), 'is heterozygote for S2S4, but homozygote for S2.'))
			else if (sample.row[,'Ag_Def_S2_4'] == 'AT')
				kdr.S.origins <- paste(kdr.S.origins, 'S2_het', sep = ',')
			else if (sample.row[,'Ag_Def_S2_4'] == 'TT')
				kdr.S.origins <- paste(kdr.S.origins, 'S4_het', sep = ',')
		}
		#
		if (is.na(sample.row[,'Ag_Def_S3']))
			kdr.S.origins <- paste(kdr.S.origins, 'S3?', sep = ',')
		else if (sample.row[,'Ag_Def_S3'] == 'GG')
			kdr.S.origins <- paste(kdr.S.origins, 'S3_hom', sep = ',')
		else if (sample.row[,'Ag_Def_S3'] == 'GT')
			kdr.S.origins <- paste(kdr.S.origins, 'S3_het', sep = ',')
		# 
		if (alternate_S4S5){
			if (is.na(sample.row[,'Ag_Def_S4S5_2']))
				kdr.S.origins <- paste(kdr.S.origins, 'S4S5?', sep = ',')
			else if (sample.row[,'Ag_Def_S4S5_2'] == 'TT'){
				if (is.na(sample.row[,'Ag_Def_S5']))
					kdr.S.origins <- paste(kdr.S.origins, '(S4S5)_hom', sep = ',')
				else if (sample.row[,'Ag_Def_S5'] == 'CC')
					kdr.S.origins <- paste(kdr.S.origins, 'S5_hom', sep = ',')
				else if (sample.row[,'Ag_Def_S5'] == 'AC')
					kdr.S.origins <- paste(kdr.S.origins, 'S5_het,S4_het', sep = ',')
				else if (sample.row[,'Ag_Def_S5'] == 'AA')
					kdr.S.origins <- paste(kdr.S.origins, 'S4_hom', sep = ',')
			}
			else if (sample.row[,'Ag_Def_S4S5_2'] == 'GT'){
				if (is.na(sample.row[,'Ag_Def_S5']))
					kdr.S.origins <- paste(kdr.S.origins, '(S4S5)_het', sep = ',')
				else if (sample.row[,'Ag_Def_S5'] == 'CC')
					return(paste('Fail. Genotypes suggest that sample', rownames(sample.row), 'is heterozygote for S4S5, but homozygote for S5.'))
				else if (sample.row[,'Ag_Def_S5'] == 'AC')
					kdr.S.origins <- paste(kdr.S.origins, 'S5_het', sep = ',')
				else if (sample.row[,'Ag_Def_S5'] == 'AA')
					kdr.S.origins <- paste(kdr.S.origins, 'S4_het', sep = ',')
			}
		}
		else{
			if (is.na(sample.row[,'Ag_Def_S4S5']))
				kdr.S.origins <- paste(kdr.S.origins, 'S4S5?', sep = ',')
			else if (sample.row[,'Ag_Def_S4S5'] == 'CC'){
				if (is.na(sample.row[,'Ag_Def_S5']))
					kdr.S.origins <- paste(kdr.S.origins, '(S4S5)_hom', sep = ',')
				else if (sample.row[,'Ag_Def_S5'] == 'CC')
					kdr.S.origins <- paste(kdr.S.origins, 'S5_hom', sep = ',')
				else if (sample.row[,'Ag_Def_S5'] == 'AC')
					kdr.S.origins <- paste(kdr.S.origins, 'S4_het,S5_het', sep = ',')
				else if (sample.row[,'Ag_Def_S5'] == 'AA')
					kdr.S.origins <- paste(kdr.S.origins, 'S4_hom', sep = ',')
			}
			else if (sample.row[,'Ag_Def_S4S5'] == 'CT'){
				if (is.na(sample.row[,'Ag_Def_S5']))
					kdr.S.origins <- paste(kdr.S.origins, '(S4S5)_het', sep = ',')
				else if (sample.row[,'Ag_Def_S5'] == 'CC')
					return(paste('Fail. Genotypes suggest that sample', rownames(sample.row), 'is heterozygote for S4S5, but homozygote for S5.'))
				else if (sample.row[,'Ag_Def_S5'] == 'AC')
					kdr.S.origins <- paste(kdr.S.origins, 'S5_het', sep = ',')
				else if (sample.row[,'Ag_Def_S5'] == 'AA')
					kdr.S.origins <- paste(kdr.S.origins, 'S4_het', sep = ',')
			}
		}
	}
	# for S heterozygotes
	else if (kdr.S.origins == 'S:het'){
		if (is.na(sample.row[,'Ag_Def_S1_3']))
			kdr.S.origins <- paste(kdr.S.origins, 'S1?', sep = ',')
		else if (sample.row[,'Ag_Def_S1_3'] == 'CC')
			return(paste('Fail. Genotypes suggest that sample', rownames(sample.row), 'is heterozygote for S kdr, but homozygote for S1.'))
		else if (sample.row[,'Ag_Def_S1_3'] == 'CT')
			kdr.S.origins <- paste(kdr.S.origins, 'S1_het', sep = ',')
		#
		if (is.na(sample.row[,'Ag_Def_S2S4']))
			kdr.S.origins <- paste(kdr.S.origins, 'S2S4?', sep = ',')
		else if (sample.row[,'Ag_Def_S2S4'] == 'TT')
			return(paste('Fail. Genotypes suggest that sample', rownames(sample.row), 'is heterozygote for S kdr, but homozygote for S2S4.'))
		else if (sample.row[,'Ag_Def_S2S4'] == 'CT'){
			if (is.na(sample.row[,'Ag_Def_S2_4']))
				kdr.S.origins <- paste(kdr.S.origins, '(S2S4)_het', sep = ',')
			else if (sample.row[,'Ag_Def_S2_4'] == 'AA')
				return(paste('Fail. Genotypes suggest that sample', rownames(sample.row), 'is heterozygote for S kdr and S2S4, but homozygote for S2.'))
			else if (sample.row[,'Ag_Def_S2_4'] == 'AT')
				kdr.S.origins <- paste(kdr.S.origins, 'S2_het', sep = ',')
			else if (sample.row[,'Ag_Def_S2_4'] == 'TT')
				kdr.S.origins <- paste(kdr.S.origins, 'S4_het', sep = ',')
		}
		#
		if (is.na(sample.row[,'Ag_Def_S3']))
			kdr.S.origins <- paste(kdr.S.origins, 'S3?', sep = ',')
		else if (sample.row[,'Ag_Def_S3'] == 'GG')
			return(paste('Fail. Genotypes suggest that sample', rownames(sample.row), 'is heterozygote for S kdr, but homozygote for S3.'))
		else if (sample.row[,'Ag_Def_S3'] == 'GT')
			kdr.S.origins <- paste(kdr.S.origins, 'S3_het', sep = ',')
		# 
		if (alternate_S4S5){
			if (is.na(sample.row[,'Ag_Def_S4S5_2']))
				kdr.S.origins <- paste(kdr.S.origins, 'S4S5?', sep = ',')
			else if (sample.row[,'Ag_Def_S4S5_2'] == 'TT'){
				return(paste('Fail. Genotypes suggest that sample', rownames(sample.row), 'is heterozygote for S kdr, but homozygote for S4S5.'))
			}
			else if (sample.row[,'Ag_Def_S4S5_2'] == 'GT'){
				if (is.na(sample.row[,'Ag_Def_S5']))
					kdr.S.origins <- paste(kdr.S.origins, '(S4S5)_het', sep = ',')
				else if (sample.row[,'Ag_Def_S5'] == 'CC')
					return(paste('Fail. Genotypes suggest that sample', rownames(sample.row), 'is heterozygote for S kdr and S4S5, but homozygote for S5.'))
				else if (sample.row[,'Ag_Def_S5'] == 'AC')
					kdr.S.origins <- paste(kdr.S.origins, 'S5_het', sep = ',')
				else if (sample.row[,'Ag_Def_S5'] == 'AA')
					kdr.S.origins <- paste(kdr.S.origins, 'S4_het', sep = ',')
			}
		}
		else{
			if (is.na(sample.row[,'Ag_Def_S4S5']))
				kdr.S.origins <- paste(kdr.S.origins, 'S4S5?', sep = ',')
			else if (sample.row[,'Ag_Def_S4S5'] == 'CC'){
				return(paste('Fail. Genotypes suggest that sample', rownames(sample.row), 'is heterozygote for S kdr, but homozygote for S4S5.'))
			}
			else if (sample.row[,'Ag_Def_S4S5'] == 'CT'){
				if (is.na(sample.row[,'Ag_Def_S5']))
					kdr.S.origins <- paste(kdr.S.origins, '(S4S5)_het', sep = ',')
				else if (sample.row[,'Ag_Def_S5'] == 'CC')
					return(paste('Fail. Genotypes suggest that sample', rownames(sample.row), 'is heterozygote for S kdr and S4S5, but homozygote for S5.'))
				else if (sample.row[,'Ag_Def_S5'] == 'AC')
					kdr.S.origins <- paste(kdr.S.origins, 'S5_het', sep = ',')
				else if (sample.row[,'Ag_Def_S5'] == 'AA')
					kdr.S.origins <- paste(kdr.S.origins, 'S4_het', sep = ',')
			}
		}
	}
	# Because both S2S4 and S4S5 markers can conclude that S4 is present, the S4_het or S4_hom output could 
	# be present twice. We therefore collapse this repeated output
	kdr.S.origins <- sub('S4_het,S4_het', 'S4_het', kdr.S.origins)
	kdr.S.origins <- sub('S4_hom,S4_hom', 'S4_hom', kdr.S.origins)
	kdr.S.origins
}

# Run the generalised linear modelling process to determine the association bewteen genetic markers and 
# insecticide resistance.
glmodelling <- function(input.table, list.of.markers = markers, rescolumn = 'AliveDead', control.for = character(), glm.function = NULL, verbose = T){
	# Check whether the markers and random effects are present in the data.frame
	if (sum(list.of.markers %in% colnames(input.table)) != length(list.of.markers))
		stop('Some of the requested markers were not found in genotypes table.')
	if (sum(control.for %in% colnames(input.table)) != length(control.for))
		stop('At least one random effect was not found in genotypes table.')
	if (!(rescolumn %in% colnames(input.table)))
		stop('Resistance column not found in genotypes table.')
	# Remove any requested random effects that have only 1 level
	random.effects <- character()
	for (this.control in control.for){
		level.counts <- tapply(input.table[,this.control], input.table[,this.control], length)
		if (max(level.counts, na.rm = T) < sum(level.counts, na.rm = T)) 
			random.effects <- c(random.effects, this.control)
		else
			cat('Removing random effect ', this.control, ' was removed because it is invariable.')
	}
	# If a glm.function was not set, decide which glm function you are going to use, based on whether mixed 
	# modelling will be necessary
	if (is.null(glm.function))
		glm.function <- ifelse(length(random.effects) > 0, 'glmer', 'glm')
	if (glm.function == 'glm'){
		# Create the random effect string, which will be empty if we have no random effects
		random.effects.string <- ifelse(length(random.effects) > 0, paste(' +', paste(random.effects, collapse = ' + ', sep = '')), '')
		# Set the name of the column containing the P.value in the anova function (which is different depending
		# on the glm function that is used)
		P.val.column <- 'Pr(>Chi)'
	}
	else if (glm.function == 'glmer'){
		random.effects.string <- ifelse(length(random.effects) > 0, paste(' +', paste('(1|', random.effects, ')', collapse = ' + ', sep = '')), '')
		P.val.column <- 'Pr(>Chisq)'
	}
	if (verbose)
		cat('Using the following string to control for confounding factors: ', random.effects.string, '\n', sep = '')
	# We remove markers for which there is no variation in the dataset or for which some alleles are too rare. 
	if (verbose)
		cat('\nDetecting invariable markers.\n')
	kept.markers <- character()
	invariable.markers <- character()
	for (this.marker in list.of.markers){
		allele.counts <- tapply(input.table[,this.marker], input.table[,this.marker], length)
		if (max(allele.counts, na.rm = T) <= (sum(allele.counts, na.rm = T) - 2)) 
			kept.markers <- c(kept.markers, this.marker)
		else
			invariable.markers <- c(invariable.markers, this.marker)
	}
	if (length(kept.markers) == 0)
		stop('Fail. None of the markers provided were variable.')
	# We do the glm analysis directly on the table from the global environment rather than the argument, this 
	# way the name of the table that was used is recorded in the modelling output. 
	working.table.name <- make.names(deparse(substitute(input.table)))
	
	# For each marker, we calculate its pseudo-R2 and P-value compared to the null model.
	if (verbose)
		cat('\nAnalysing markers independently.\n')
	individual.markers <- data.frame(P = numeric(), pseudo.R2 = numeric())
	for (this.marker in kept.markers){
		# Remove the Na values for this marker
		this.table <- input.table[!is.na(input.table[,this.marker]),]
		# Build the model 
		this.model <- eval(parse(text = paste(glm.function, '(', rescolumn, ' ~ ', this.marker, random.effects.string, ', data = this.table, family = binomial)', sep = '')))
		# Build the null model
		this.null.model <- eval(parse(text = paste(glm.function, '(', rescolumn, ' ~ 1 ', random.effects.string, ', data = this.table, family = binomial)', sep = '')))
		# Get the stats
		this.p <- anova(this.model, this.null.model, test = 'Chisq')[[P.val.column]][2]
		# Report pseudo Rsquared if we used GLM and if we have the modEvA package
		if (('modEvA' %in% (.packages())) & (glm.function == 'glm'))
			this.pseudo.r2 <- mean(unlist(RsqGLM(this.model)))
		else
			this.pseudo.r2 <- NA
		individual.markers[this.marker, ] <- c(this.p, this.pseudo.r2)
	}
	
	# We now build the full model and remove markers one by one until all markers are significant
	markers.remaining <- kept.markers
	full.model.text <- paste(glm.function, '(', rescolumn, ' ~ ', paste(markers.remaining, collapse = ' + '), random.effects.string, ', data = ', working.table.name, ', family = binomial)', sep = '')
	# We'll keep track of the markers with perfect correlation
	correlated.markers <- character()
	if (verbose)
		cat('\nRunning commentary on model optimisation:\n\n')
	while(length(markers.remaining)){
		# It is possible, when building the model, that some of the markers become monomorphic (because the other 
		# alleles are removed when other markers are NA). The model building will raise an error if this is the 
		# case, so let's check for this and temporarily remove such markers before we go ahead.
		temporarily.removed <- character()
		markers.remaining.genotypes <- input.table[, markers.remaining, drop = F]
		markers.remaining.genotypes <- markers.remaining.genotypes[complete.cases(markers.remaining.genotypes), , drop = F]
		for (this.marker in markers.remaining){
			allele.counts <- tapply(markers.remaining.genotypes[ ,this.marker], markers.remaining.genotypes[,this.marker], length)
			if (max(allele.counts, na.rm = T) == sum(allele.counts, na.rm = T)) {
				if (verbose){
					cat('Temporarily removing marker ', this.marker, ' as it has no variation left when NAs at other ',
					    'loci are removed.\n', sep = '')
				}
				temporarily.removed <- c(temporarily.removed, this.marker)
			}
		}
		# If any markers need to be temporarily removed, do so here.
		working.markers <- markers.remaining[!(markers.remaining %in% temporarily.removed)]
		# Build the model using the working.markers
		old.model.text <- paste(glm.function, '(', rescolumn, ' ~ ', paste(working.markers, collapse = ' + '), random.effects.string, ', data = ', working.table.name, ', family = binomial)', sep = '')
		if (verbose)
			cat('Building model:', old.model.text, '\n')
		old.model <- eval(parse(text = old.model.text))
		p.values <- numeric()
		# This will tell us whether we have broken out of the marker loop and therefore should go to the next model loop
		next.loop <- F
		for (this.marker in working.markers){
			new.model <- eval(parse(text = paste('update(old.model, .~.-', this.marker, ')', sep = '')))
			# Check that the new model doesn't have more rows than the old one (if the removed marker had unique NAs)
			if (length(fitted(new.model)) == length(fitted(old.model))){
				this.p.value <- anova(old.model, new.model, test = 'Chisq')[[P.val.column]][2]
			}
			# Otherwise, we need to rebuild the models with those samples removed.
			else{
				reduced.input.table <- input.table[!is.na(input.table[,this.marker]),]
				temp.old.model <- eval(parse(text = paste(glm.function, '(', rescolumn, ' ~ ', paste(working.markers, collapse = ' + '), random.effects.string, ', data = reduced.input.table, family = binomial)', sep = '')))
				new.model <- eval(parse(text = paste('update(temp.old.model, .~.-', this.marker, ')', sep = '')))
				this.p.value <- anova(temp.old.model, new.model, test = 'Chisq')[[P.val.column]][2]
			}
			# If the p.value was NA, then there is perfect correlation or something else was wrong. Remove the marker 
			# with a comment
			if (is.na(this.p.value)){
				if (verbose)
					cat('\tRemoving marker ', this.marker, ' due to perfect correlation.\n\n', sep = '')
				markers.remaining <- markers.remaining[markers.remaining != this.marker]
				correlated.markers <- c(correlated.markers, this.marker)
				if (length(markers.remaining) == 0){
					final.model <- paste(glm.function, '(', rescolumn, ' ~ ', '1 ', random.effects.string, ', data = ', working.table.name, ', family = binomial)', sep = '')
					if (verbose)
						cat('All variables removed.\n')
				}
				next.loop <- T
				break
			}
			else{
				p.values[this.marker] <- this.p.value
			}
		}
		if (next.loop)
			next
		else if (max(p.values) > 0.05){
			# Remove the highest non-significant p-value
			if (verbose)
				cat('\tRemoving marker ', names(p.values)[which.max(p.values)], ' as the highest, non-significant marker (P = ', max(p.values), ').\n\n', sep = '')
			marker.to.remove <- names(p.values)[which.max(p.values)]
			markers.remaining <- markers.remaining[markers.remaining != marker.to.remove]
			if (length(markers.remaining) == 0){
				# If there are no markers remaining, then the final model is the null model
				final.model <- eval(parse(text = paste(glm.function, '(', rescolumn, ' ~ ', '1 ', random.effects.string, ', data = ', working.table.name, ', family = binomial)', sep = '')))
				if (verbose)
					cat('\tAll variables removed.\n')
			}
		}
		else {
			final.model <- old.model
			if (verbose){
				cat('\tAll remaining variables significant, keeping final model:\n')
				print(final.model)
			}
			break
		}
	}
	if (verbose){
		if (length(markers.remaining) == 0)
			cat('Final model was the null model.\n\n')
		else {
			cat('Final model contained ', length(markers.remaining), ' parameters: ', paste(markers.remaining, collapse = ','), '.\n\n', sep = '')
			if (length(temporarily.removed) > 0){
				cat('\tMarkers ', paste(temporarily.removed, collapse = ','), ' were temporarily removed but could ',
					'never be reintegrated into the model.\n', sep = '')
			}
		}
	}
	# Now get the p-values and pseudo R-squared value for all the variables in the final model, when added as the 
	# last variable
	if (length(markers.remaining) > 0){
		deviance.effect <- numeric()
		for (this.marker in names(p.values)){
			reduced.model <- eval(parse(text = paste('update(final.model, .~.-', this.marker, ')', sep = '')))
			deviance.effect[this.marker] <- deviance(reduced.model) - deviance(final.model)
		}
	}
	else {
		p.values <- NA
		deviance.effect <- NA
		temporarily.removed <- NA
	}
	cat('\n')
	#
	final.model.sig <- data.frame(P = p.values, deviance = deviance.effect)
	# Return the output
	list('invariable.markers' = invariable.markers, 'correlated.markers' = correlated.markers, 'sig.alone' = individual.markers, 'final.model' = final.model, 'final.sig' = final.model.sig, 'temporarily.removed' = temporarily.removed)
}
